# Distribution and functional potential of photoautotrophic bacteria in alkaline hot springs

**DOI:** 10.1101/2020.12.16.423123

**Authors:** Annastacia C. Bennett, Senthil K. Murugapiran, Eric D. Kees, Trinity L. Hamilton

## Abstract

Alkaline hot springs in Yellowstone National Park (YNP) provide a framework to study the relationship between photoautotrophs and temperature. Previous work has focused on cyanobacteria (oxygenic phototrophs), but anoxygenic phototrophs are critical parts of the evolutionary history of life on Earth and and are abundant across temperature gradients in alkaline hot springs. However, many questions remain regarding the ecophysiology of anoxygenic photosynthesis due to the taxonomic and metabolic diversity of these taxa. Here, we examined the distribution of genes involved in phototrophy and carbon and nitrogen fixation in eight alkaline (pH 7.3-9.4) hot spring sites approaching the upper temperature limit of photosynthesis (~72°C) in YNP using metagenome sequencing. Genes associated with cyanobacteria are abundant throughout our data and more diverse at temperatures > 63°C, genes for autotrophic Chloroflexi are more abundant in sites > 63°C and genes associated with phototrophic Chloroflexi are abundant throughout. Additionally, we recovered deep branching nitrogen fixation genes from our metagenomes, which could inform the evolutionary history of nitrogen fixation. Lastly, we recovered 25 metagenome assembled genomes of Chloroflexi. We found distinct differences in carbon fixation genes in *Roseiflexus* and *Chloroflexus* bins, in addition to several novel Chloroflexi bins. Our results highlight the physiological diversity and evolutionary history of the understudied, anoxygenic autotrophic Chloroflex. Furthermore, we provide evidence that genes involved in nitrogen fixation in Chloroflexi is more widespread than previously assumed.

**IMPORTANCE:** Photosynthetic bacteria in hot springs are of great importance to both microbial evolution and ecology because they are responsible for the rise of oxygen and are critical to nutrient cycling. While a large body of work has focused on the oxygenic photosynthesis in cyanobacteria, many questions remain regarding the metabolic potential of anoxygenic phototrophs but are further compounded by their metabolic and taxonomic diversity. Here, we have recovered several novel metagenome bins and quantified the distribution of key genes involved in carbon and nitrogen metabolism in both oxygenic and anoxygenic phototrophs. Together, our results add to the body of work focusing on photosynthetic bacteria in hot springs in Yellowstone National Park.

## INTRODUCTION

A rich history of research on microbial communities in hot springs in Yellowstone National Park (YNP) has revealed that photoautotrophic bacteria are the main primary producers in alkaline hot springs > 60°C to 72°C (reviewed in 1). Alkaline hot springs > ~56°C are typically devoid of eukaryotic life, contain fine-scale temperature gradients and, therefore, are excellent environments to study the ecophysiology of photoautotrophs *in situ* (2). Historically, cyanobacteria have been the focus of these studies as they are the only bacteria capable of oxygenic photosynthesis—a process that arose >2.2 billion years ago and is of great importance to Earth’s history (3). The distribution of cyanobacterial species in Mushroom and Octopus Springs, two alkaline hot springs in YNP, has been described at ~ 1°C resolution (4–7), revealing ecotypes that are highly adapted to specific temperatures. Additionally, cyanobacterial diversity increases with decreasing temperature (8). While anoxygenic phototrophs are also present in high temperature alkaline springs (5,6,8–10), many questions remain regarding their metabolic potential and contribution to carbon and nitrogen cycling *in situ,* which is complicated by their taxonomic and metabolic diversity.

While Cyanobacteria have been widely studied for several decades, researchers have only begun to ascertain the metabolic potential of phototrophic Chloroflexi. In alkaline, high temperature hot springs, cyanobacteria are the only organisms that perform oxygenic photosynthesis, using two photosystems in concert to harvest electrons from water for metabolic processes like carbon fixation. In contrast to oxygenic phototrophs, anoxygenic phototrophs require one light-harvesting reaction center complex – either type I or type II – and have been observed in seven additional bacterial phyla that persist in alkaline hot springs (1). Of these phyla, Chloroflexi are the most abundant anoxygenic phototrophs in alkaline hot springs > 60°C, while other anoxygenic phototrophs (e.g. Chlorobi and *Candidatus* Chloracidobacteium thermophilum), are usually present but less abundant (8,9,11,12). At present, phototrophy in hot spring Chloroflexi is limited to class *Chloroflexales,* with the exception of the novel photoheterotroph, *Candidatus* Roseilineae in class Anaerolineae (13). To date, there have been a handful of isolate and *in situ* studies on *Chloroflexus* and *Roseiflexus.* For example, in pure culture, *Roseiflexus castenholzii* has not been grown in the absence of fixed carbon or nitrogen, yet *Roseiflexus* genomes recovered from alkaline hot springs contain genes involved in both of these processes (14). Additionally, we observe *Chloroflexus* in alkaline hot springs up to ~70°C; however, in isolate studies, carbon fixation using the 3-hydroxypropionate bicycle (3HPB) by *Chloroflexus* has not been demonstrated at temperatures > 55°C (15). Therefore, many questions remain regarding the distribution and metabolic potential of the phototrophs in class *Chloroflexales* and of novel classes within the phylum.

Photosynthesis is essential to cycle carbon in hot springs. However, these environments are also nitrogen limited, which allows nitrogen-fixing bacteria to thrive in hot springs. Because oxygen solubility decreases as a function of increased temperature, oxygen concentrations are often low, creating conditions that are suitable for the production of the oxygen-sensitive enzyme, nitrogenase (16,17). Nitrogenase is an iron-sulfur complex containing one of three metals harbored in the active site: molybdenum (Mo), iron (Fe) or vanadium (V). All three varieties exist in nitrogen limited environments, but Mo-nitrogenase is the most common and is encoded by *nif* genes (18,19). *nif* genes are dispersed throughout environments on Earth, especially areas that are typically nitrogen-limited, including alkaline hot springs in YNP (20,21). In-depth studies quantifying the distribution of *nifH* (which encodes the iron protein (NifH) in nitrogenase) and potential nitrogenase activity have been conducted in a number of acidic hot springs > 55°C in YNP (22,23). These studies revealed that diazotrophs in acidic hot springs are highly adapted to local conditions. In alkaline hot springs, transcriptional activity of *nifH* in *Leptococcus* (Cyanobacteria, *Synechococcus* renamed to *Leptococcus* in (24)) species was monitored over a diel cycle in 53-63°C mats which revealed an increase in activity at the end of the day, once mats turned anoxic (16,25). However, broader studies linking the distribution of *nifH* to phototrophic community composition have not been conducted in alkaline springs > 63°C, to our knowledge. Additionally, *Roseiflexus* genomes contain *nif* genes, but they lack the full protein suite required to build a functional nitrogenase and likely do not fix carbon *in situ,* but the functional purpose of NifH in *Roseiflexus* remains unknown.

Previous work in YNP has largely relied on single marker gene or metagenome assembled genome (MAG, herein referred to as ‘bin’) abundance to define the range of photosynthesis in alkaline hot springs *in situ*(1–4). Our previous work suggested that Chloroflexi and cyanobacteria were highly abundant in alkaline hot springs ranging from 60°C to 72°C (12). Given the abundance of cyanobacteria and Chloroflexi in these sites and the crucial role that they play in nitrogen and carbon cycling, we sought to determine the distribution and functional potential of phototrophic bacteria in a subset of hot springs from Hamilton *et al.* 2019. Here, we overcame the limitations of primer bias in single gene surveys and potentially contaminated/incomplete metagenome bins and examined genes and pathways associated with phototrophy and nitrogen fixation in metagenomes as has been proven informative in other systems (23). We sequenced metagenomes from eight alkaline hot spring samples in YNP with temperatures ranging from 62°C to 71°C (Table 1). We found that 1) genes associated with cyanobacterial photoautotrophy are abundant throughout our data and more diverse at temperatures > 63°C, 2) genes for autotrophic Chloroflexi are more abundant in sites > 63°C, 3) genes associated with phototrophic Chloroflexi are abundant throughout, and 4) we recovered deep branching nitrogen fixation genes from our metagenomes. Lastly, we binned our metagenome assemblies and recovered 25 Chloroflexi bins. We found distinct differences in carbon fixation genes in *Roseiflexus* and *Chloroflexus* bins, in addition to several novel Chloroflexi bins. Together, these results add to the body of work on photoautotrophic bacteria in alkaline hot springs which is critical to solving the evolutionary history and ecophysiology of nitrogen fixation and photosynthesis in bacteria.

**Table 1.**
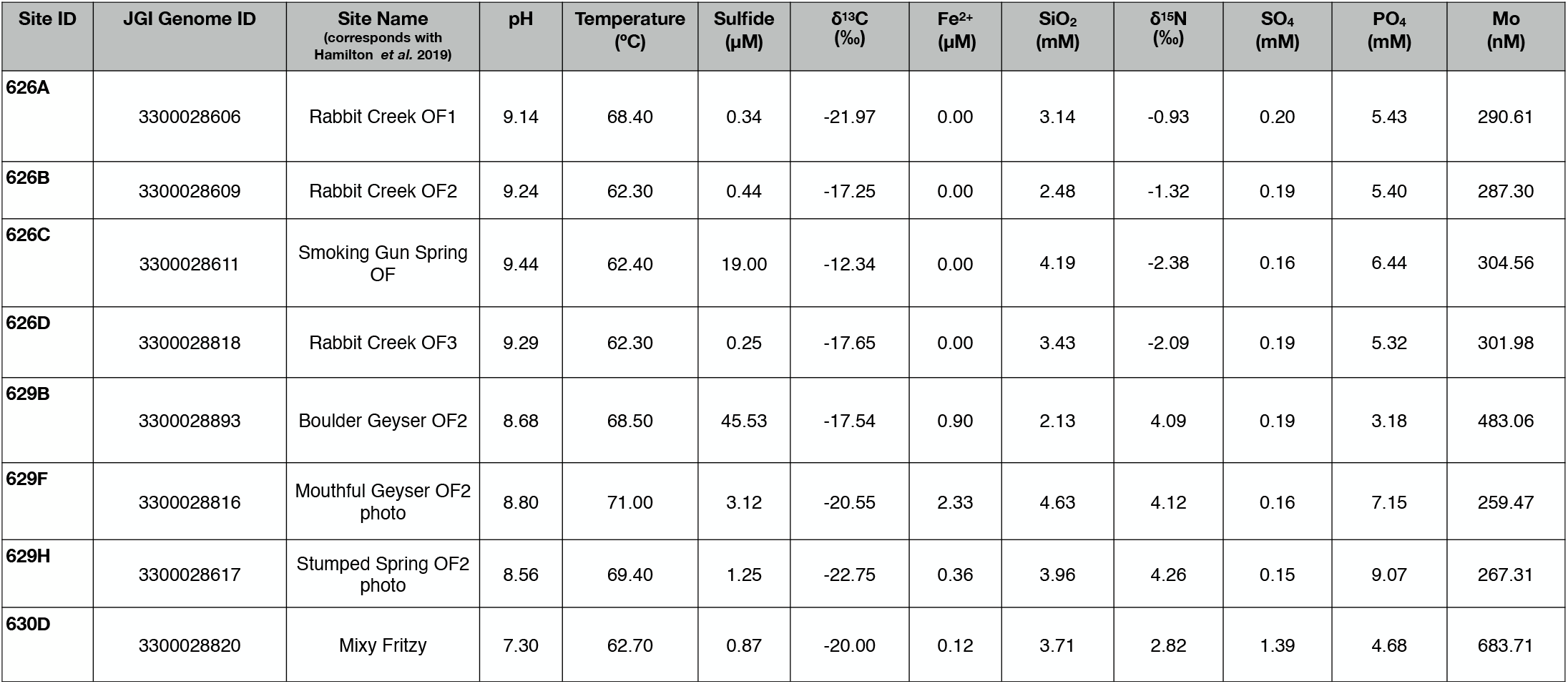
Geochemistry of sites.

## Results and Discussion

### Overview of site geochemistry and metagenome assemblies

In Hamilton *et al.* 2019 (12), we surveyed the distribution and putative activity of phototrophy in 22 hot springs using 16S rRNA gene sequencing, ^13^C and ^15^N isotopic signatures and photosynthetic microcosm experiments. We report that a large proportion of the population in sites > 60°C and pH > 7 are putative phototrophic Chloroflexi and Cyanobacteria, namely *Leptococcus, Roseiflexus* and *Chloroflexus*. Our 2019 study, in addition to several other studies, suggest that Cyanobacteria and phototrophic Chloroflexi are the dominant photoautotrophs in alkaline hot springs from 60°C to the upper temperature limit of photosynthesis (72°C) (1,5,14,28,29). To this end, we generated metagenome sequences from a subset of samples (n=8, Table 1) to determine the distribution of specific genes involved in photoautotrophy and nitrogen fixation in YNP alkaline hot springs that approach the upper temperature limit of photosynthesis. The sites range in temperature from 62°C to 71°C and pH between 7 and 9 (Table 1, (12)). The average number of reads between the eight metagenomes was 830473, with a standard deviation of 267811 reads (Table S1). In our metagenome assemblies, the maximum number of reads was from site 629F (1187870 reads) while the lowest number of reads was from 626C (375420 reads) (Table S1). Site 626B contained the highest number of open reading frames, 332336 (ORFs, e.g. genes), while 626C had the lowest number of genes, 150190 (Table S1).

### Photosynthetic reaction center alpha diversity varies with temperature

Oxygenic photosynthesis is a remarkable metabolism that involves two photosystems, Photosystem I (PSI) and Photosystem II (PSII), working in concert to harvest electrons from water to fuel carbon fixation and other cellular processes. PSII houses the oxygen evolving complex where light energy is captured to liberate electrons from water — a process that requires manipulation of five proteins (encoded by *psb* genes) surrounding the oxygen evolving complex (30). In our data, we quantified the abundance of three *psb* genes: *psbA*, *psbB* and *psbD. psbA* and *psbD* encode for the D1 and D2 proteins, respectively, which both serve to ligate the redox active components in PSII and are highly transcriptionally regulated in Cyanobacteria (31,32). *psbB* encodes CP47, a chlorophyll binding protein crucial to forming a stable PSII reaction center (33). In our metagenomes, abundance of *psb* genes varied between sites (Figure 1). We observed larger mean abundances of *psb* genes at temperatures above 68°C and the highest mean abundance of *psb* genes occurred in the site with the highest temperature (71°C, 629F, Figure 1), while the lowest mean abundance was observed at 69.4°C (site 629H), suggesting other factors limit cyanobacteria in 629H. However, in sites < 63°C, we recovered more copies of *psb* genes, suggesting higher alpha diversity within the cyanobacteria population or that single species harbor multiple copies of the *psb* genes. Previous work has shown that cyanobacterial diversity in alkaline hot springs decreases with increasing temperature and that *Leptococcus* species, which contain up to three copies of *psbA* (4) are most abundant in these conditions (8,9,12).

**Figure 1.**
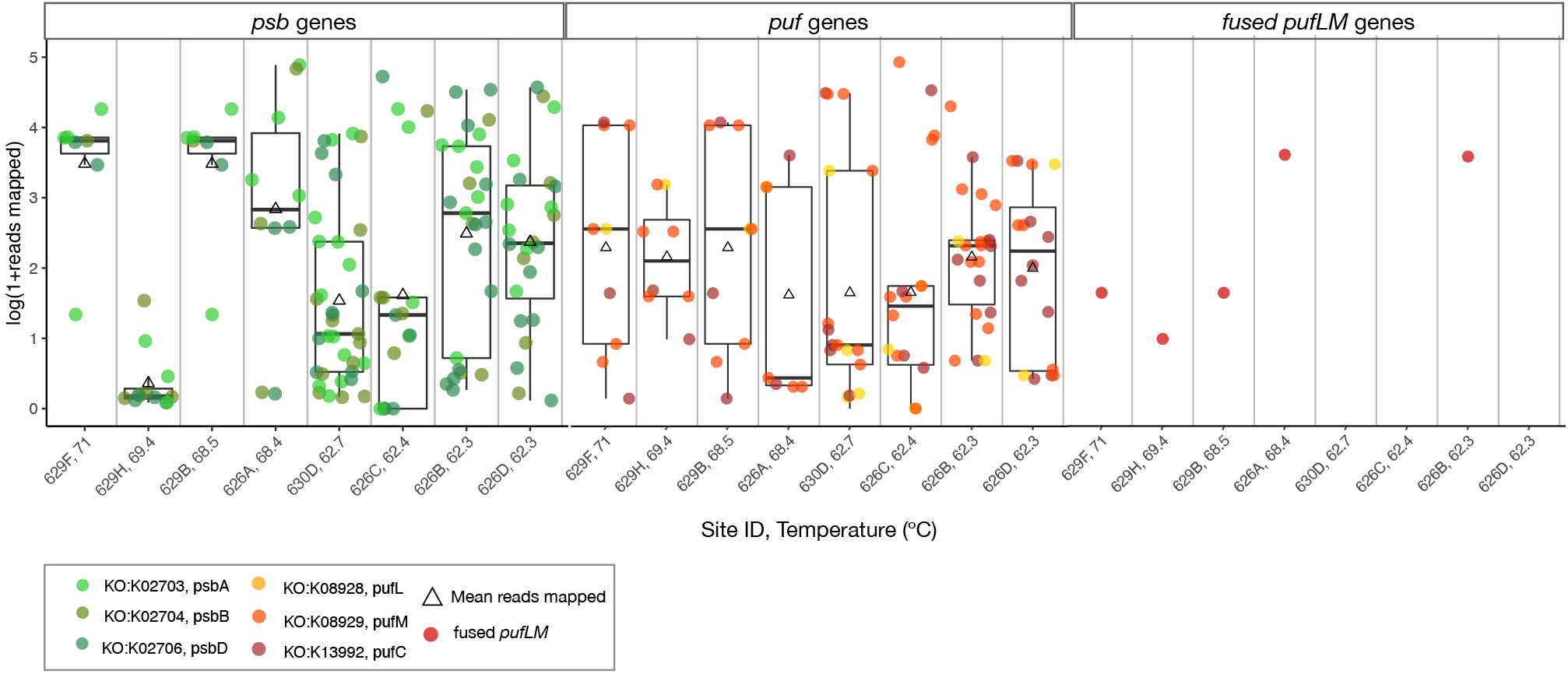
Distribution of photosynthetic machinery with temperature. The overall abundance (natural log of 1 + reads mapped) of genes that encode for Cyanobacterial photosystem II (*psb*) and type II anoxygenic photosynthesis reaction centers (*puf* and fused *pufLM*) are shown as box plots for each site. Triangles represent the mean abundance for the gene set and dots represent individual gene abundances, shaded by the corresponding photosystem or reaction center gene. Box plot outliers were removed. Sites are ordered by decreasing temperature.

In all organisms that perform oxygenic photosynthesis, the *psbA* gene encodes the D1 protein — a core protein of photosystem II. Several cyanobacteria genomes contain multiple copies and variant forms of the *psbA* gene to rapidly repair the D1 protein or invoke a variant isoform in response to environmental conditions (32). Typically, cyanobacteria encode a ‘standard’ form (usually *psbA1*), expressed under normal conditions, and a ‘stress-induced’ form (*psbA2-n*), expressed under high-stress conditions such as low temperature, high UV irradiance or microaerobic conditions (31). For example, in marine cyanobacteria, the stress-induced isoform (encoded by *psbA2*) is preferentially expressed under high UV conditions (34). Fine-scale sampling of *Leptococcus* in Octopus Spring revealed there are five *Leptococcus* ecotypes (A′′, A′, A, B′, and B, in order of decreasing thermotolerance) that persist between 73°C to 50°C and are adapted to specific temperatures (7,35). Reference sequences from an A-type and B’-type ecotype – *Leptococcus yellowstonii* JA-3-3Ab and *Leptococcus springii* JA-2-3B’a(2-13) respectively (35)– were used to identify *psbA* genes in our samples. Henceforth, we will refer to the *Leptococcus* A and B ‘ecotypes, which respectively occupy temperature ranges of ~55-68°C and ~50-62°C, as either A-type or B’-type. Each of the YNP *Leptococcus* genomes contain two to three variants of the *psbA* gene, but the environmental conditions that select for strains that harbor multiple copies is unknown.

In the present study, we recovered several *Leptococcus-*like PsbA sequences. To determine whether they were most closely related to the A- or B’-type *Leptococcus* PsbA variants, we built a phylogenetic tree with our *Leptococcus*-like PsbA sequences and both A-type and B’-type reference sequences (Figure 2). 17 of the 22 PsbA sequences in our data were most closely related to PsbA from one of the two *Leptococcus* reference strains and, in general, we did not observe a preference for either PsbA1 or PsbA2 with temperature. The B’-type PsbA2 and A-type PsbA1 references formed a separate clade containing six of our PsbA sequences. Bifurcating in a deeper branch from this clade, the A-type PsbA2 and B’-type PsbA1 references grouped with the remaining eight *Leptococccus* PsbA sequences. Within each of those clades, several sequences branch outside of the PsbA1 and PsbA2 reference sequences, suggesting a third PsbA sequence is present in our metagenomes.

**Figure 2.**
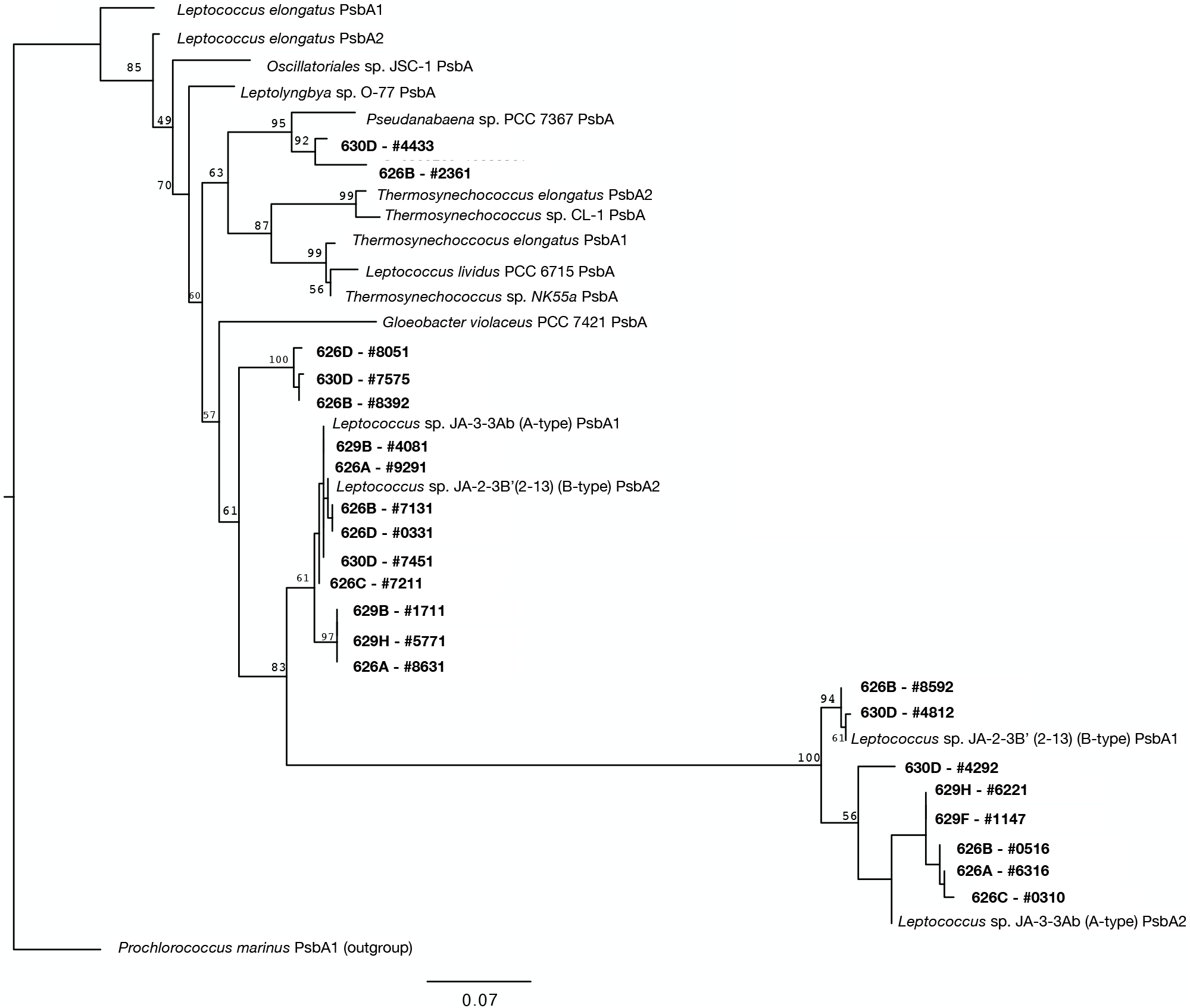
Maximum likelihoodinferred PsbA tree. Maximum likelihood-inferred phylogenetic reconstruction of all PsbA protein sequences recovered from YNP sites (22 sequences) (bold) aligned to references sequences. The tree is rooted with a marine Cyanobacterium, *Prochlorococcus marinus* PsbA1, as an out group. The scale bar indicates less than 1 substitution for every ten positions. Bootstrap values (n=1000) are displayed at branches. Bootstrap values less than 40 are not shown.

Anoxygenic photosynthesis requires only one light-harvesting reaction center complex—type I or type II. *psc* genes encode Type I reaction centers (in Chlorobi, Firmicutes *(Heliobacteriaceae)*, Acidobacteria (*Candidatus* Chloracidobacteium thermophilum)), while *puf* genes encode type-II reaction centers (in Chloroflexi, (class *Chlorofexales,* and *Candidatus* Thermofonsia), Proteobacteria *(Alpha-, Beta-, and Gamma-)* and Gemmatimonadetes (strain AP64)) (1,36–38). In our metagenomes, *psc* genes were more abundant in sites below 63°C while *puf* genes were abundant in all sites (Figure 1, Figure S1). Similar to *psb* genes, we recovered more copies of both *psc* and *puf* genes in sites < 63°C, suggesting diversity (or taxa) abundance increases with decreasing temperature. These results are consistent with our previous work that suggest phototrophs with type-I RCs are less abundant at high temperatures, while the type-II phototrophic Chloroflexi are highly abundant and diverse in 60-72°C hot springs (8,12).

In anoxygenic phototrophs with type II reaction centers, genes that encode essential functions differ between species. *pufL* and *pufM* are common to all type-II phototrophs and encode PufL and PufM membrane-spanning proteins that bind bacteriochlorophylls in type-II reaction centers (39). Both *pufL* and *pufM* genes are required to form a functional reaction center yet in our data, we recovered 5:1 *pufM:pufL* genes in sites > 63°C (Figure 1). In *Roseiflexus* and *Kouleothrix* species *pufL* and *pufM* are fused into a single gene (37). We recovered five fused *pufLM* sequences in our dataset (Figure 1) that were represented in five of our eight sites. They were the most abundant in 626A and 626B, two sites in the Rabbit Creek area, where *Roseiflexus* are common and abundant (Bennett *et al.* 2020).

### Abundance and diversity of carbon cycle machinery varies with temperature

Photoautotrophic bacteria fix the majority of carbon in alkaline geothermal springs by using one of three different pathways. Cyanobacteria use the Calvin-Benson-Bassham (CBB) cycle, Chlorobi use the reductive tricarboxylic acid (rTCA) cycle and photoautotrophic Chloroflexi use the 3-hydroxypropionate bicycle (3HPB) (reviewed in (41)) with the exception of CBB-containing *Kouleothrix aurantiaca* isolated from activated sludge (42). In our data, mean reads mapped for the CBB cycle surpassed the 3HPB and rTCA in all but two sites (626C and 626B) (Figure 3B). In the CBB cycle, the carboxylation step is carried out by the enzyme ribulose 1,5 bisphosphate carboxylase/oxygenase (RuBisCO, encoded by *rbcL* [large subunit] and *rbcS* [small subunit] genes). RuBisCO, the workhorse of the CBB cycle, is highly abundant in several environments on Earth and considered to be the most abundant protein in the world (43). In hot springs, specifically, *Leptococcus* species have evolved a thermotolerant form of RuBisCO that functions at up to 74°C – toxic intermediates (reactive oxygen species) form above this temperature (44,45). Phosphoribulokinase (encoded by the *prk* gene), a second essential step of the CBB cycle, does not appear to have an upper temperature limit beyond that of phototrophy, but is likely only present in organisms that use the CBB cycle (46).

**Figure 3.**
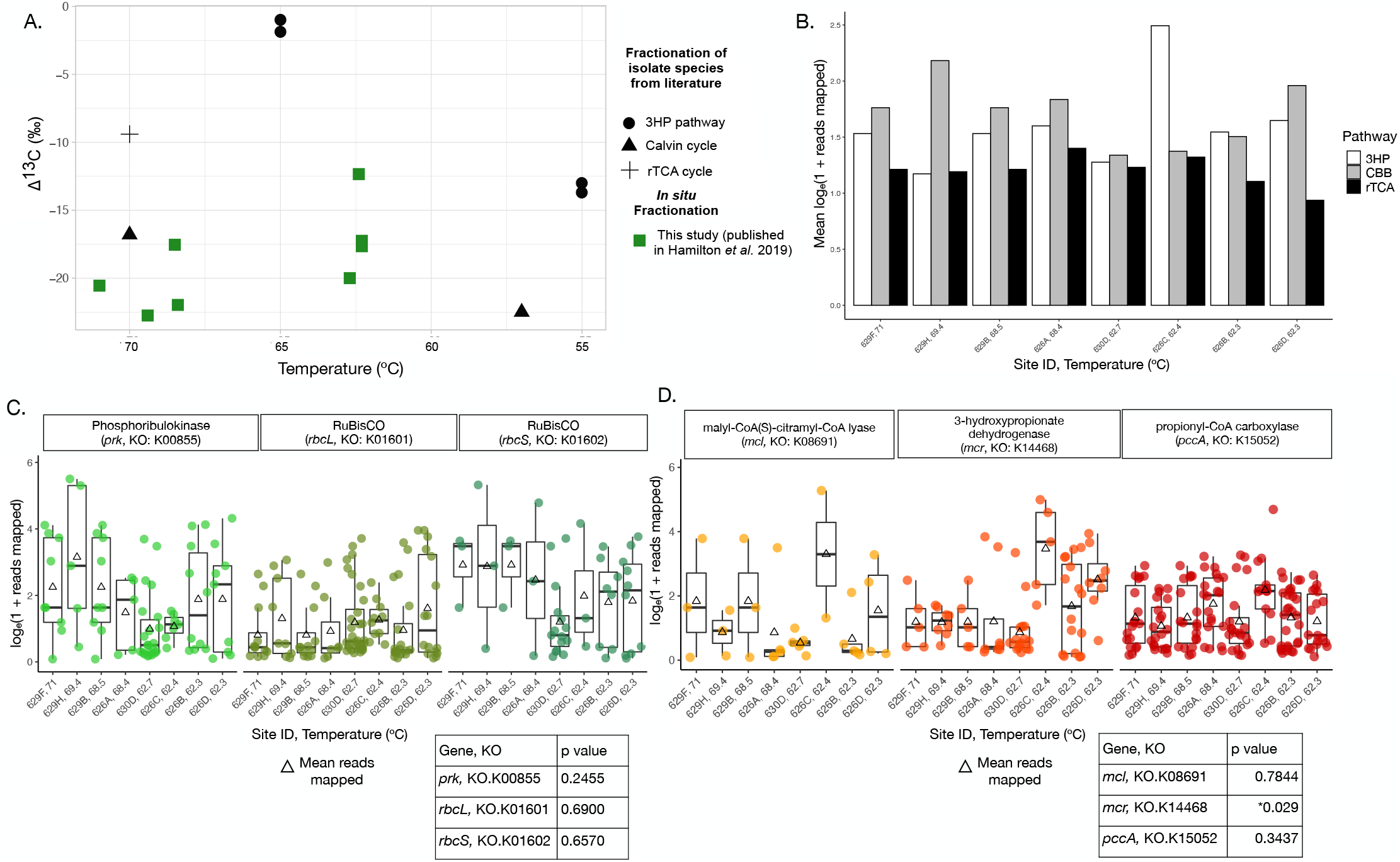
^13^C isotopic signatures and distribution of genes involved in phototrophic carbon fixation pathways. Carbon stable isotope signals of biomass are plotted by site temperature. (B) The mean abundance (natural log of 1 + reads mapped) for three photoautotrophic carbon fixation pathways are shaded by pathway. Mean abundances are calculated from three representative genes for each pathway shown in (B), the abundance (natural log of 1 + reads mapped) of gnes that encode for the Calvin cycle (C) and the 3-hydroxypropionate bicycle (D) are shown as box plots for each site. Triangles represent the mean abundance for the gene set and dots represent individual gene abundances, shaded by the genes. Box plot outliers were removed. Sites are ordered by decreasing temperature. Below each box plot are tables of Kruskal-Wallis H test significance values (p-value) for each gene.

While it is known that characteristic enzymes in the CBB cycle function at high temperatures, we sought to compare abundances of these genes across sites (Figure 3C). We observed larger mean abundances of *rbcS* than *rbcL,* but more copies of *rbcL* than *rbcS,* suggesting the CBB taxa could contain more copies of *rbcL*, or multiple forms of RuBisCO are present in these high temperature, alkaline hot springs. Furthermore, we computed the ratio of *rbcL:rbcS* and *prk:rbcS* with temperature (Figure S2) and found that both ratios were higher (more *rbcL* and *prk* genes) in sites with low temperature. At present, four forms of RuBisCO exist in nature: form I RuBisCO (cyanobacteria, alpha-, beta-, gamma-proteobacteria, chloroflexi and higher eukaryotes) is encoded by both *rbcL* and *rbcS* genes, while forms II (alpha-, beta-, gamma-proteobacteria) and III (only in methanogenic archaea) contain only the large subunit (encoded by *rbcL* genes) (47). Our data suggest that more distinct CBB taxa are present at low temperatures, or that form II or III RuBisCO taxa (*rbcL* only, non-cyanobacterial CBB cycle) persist at lower temperatures.

Chlorobi are the only phototrophic group that fix carbon via the rTCA cycle, but this pathway is distributed across several non-phototrophic lineages that are often recovered from hot springs (e.g. in the Aquificae phylum; 12,17). In general, reads associated with the rTCA cycle were low compared to other two pathways (Figure 3B). Additionally, we recovered very few reads associated with ATP citrate-lyase, an irreversible and critical enzyme in the rTCA cycle (Figure S3), suggesting few taxa fix carbon using this pathway between 62-71°C.

In contrast to the rTCA cycle, reads associated with genes involved in 3HPB, the carbon fixation pathway in most photoautotrophic Chloroflexi, were widespread and abundant in our metagenomes (Figure 3B). The 3HPB requires two carboxylation steps (via acetyl-CoA carboxylase and propionyl-CoA carboxylase), followed by steps that generate 3-hydroxypropionate and glyoxylate intermediates (48). To this end, we surveyed the abundance of three genes that are involved in these critical steps in the pathway: malyl-CoA/citramyl-CoA lyase (*mcl* gene, glyoxylate generation), propionyl-CoA carboxylase (*pccA* gene, CO2 carboxylation) and 3-hydroxypropionate dehydrogenase (*mcr* gene, 3-hydroxypropionate generation). We found that the mean abundances of *mcr* showed high variation between sites (Kruskal-Wallis H test, p < 0.05), while *pccA* and *mcl* abundances were relatively consistent across sites (Figure 3D). It was previously thought that the 3HPB pathway was only present in Archaea until *Chloroflexus* isolates were grown autotrophically using the 3HPB (48,49). While *Roseiflexus castenholzii* has not been grown without acetate (50), transcripts for the 3HPB have been recovered from metatranscriptomes of *Roseiflexus* species *in situ* (14). Previous work has shown that *Roseiflexus* and *Chloroflexus* species are abundant in alkaline hot springs >60°C and ^13^C fractionation values suggest the 3HPB could be active in these sites (8,14,20).

Years of research on primary productivity in hot springs has shown that autotrophic bacteria produce carbon isotope fractionation values that correspond to specific carbon fixation pathways (reviewed in (20)). To test our hypothesis that the CBB cycle is more prominent in samples > 68°C and the 3HPB in samples < 63°C, we compared the isotopic fractionation of ^13^C in our samples to fractionation values of characterized isolates (Figure 3A) (12). Carbon isotope fractionation in high temperature samples (>68°C) were similar to fractionation of YNP isolate *Synechococcus lividis* grown autotrophically at 70°C (51), while the low temperature samples grouped together within range of *Chloroflexus* 3HPB fractionation values at 55°C (15). Carbon isotope fractionation of isolates fixing carbon via the 3HPB have not been reported within range of our sites, but both the abundance of *puf* genes (Figure 1), 3HPB genes in our samples (Figure 3D) and ^13^C fractionation data suggest this pathway in phototrophic Chloroflexi could play an important role in primary productivity in alkaline hot springs >62°C.

### Deeply-branching nifH genes are abundant in alkaline hot springs

We quantified the distribution and abundance of *nifH* genes in eight metagenome assemblies, spanning six alkaline hot springs with temperatures from 62 to 71°C to determine if *nifH* abundances vary with temperature (Figure S4). We recovered more *nifH* genes in sites below 63°C (Figure S5B) while the mean abundance of *nifH* was highest in 626A and lowest in 629H, both sites with temperatures > 68°C. Spearman-rank correlations of our *nifH* genes with temperature, sulfide, total iron and molybdenum revealed that none of these environmental parameters correlated with abundance of *nifH* genes in our samples (Figure S4C). Together, these results suggest that *nifH* abundance does not correlate with environmental parameters, but more diverse diazotrophs are present at lower temperatures.

To determine the taxa associated with our *nifH* sequences, we constructed a phylogenetic tree with translated *nifH* gene sequences (Figure 4). Two NifH sequences were closely-related to cyanobacteria species, notably *Leptococcus* JA-3-3Ab, a common constituent of alkaline hot springs > 60°C and a known diazotroph (6,25). Four of the five most abundant NifH sequences in our dataset were closely related to *Roseiflexus* species from both high and low temperature sites (Figure 4). *nifH* genes are present in *Roseiflexus* genomes and *nifH* transcripts *Rosiefleus* have been observed *in situ* (14), but they lack the full gene suite required to build a functional nitrogenase— *Roseiflexus* genomes only encode *nifHBDK* (52) – and neither of the two isolate species (*R. castenholzii* or *Roseiflexus* sp. RS-1) can grow in the absence of fixed nitrogen (50,52), therefore, it is unlikely that *Roseiflexus* fix nitrogen. However, *Roseiflexus nifH* could be important to determining the evolutionary history of nitrogenase – *nifH* shares an evolutionary history with *bchL,* a gene involved in chlorophyll biosynthesis in anoxygenic phototrophs and BchL and NifH have high sequence and structure similarity but are functionally different (53). Lastly, while *Roseiflexus* species lack a full gene set to fix nitrogen in alkaline hot genes are abundant in our data and *nifH* mRNA has been detected in similar hot springs (14), suggesting NifH serves a functional purpose that remains unknown.

**Figure 4.**
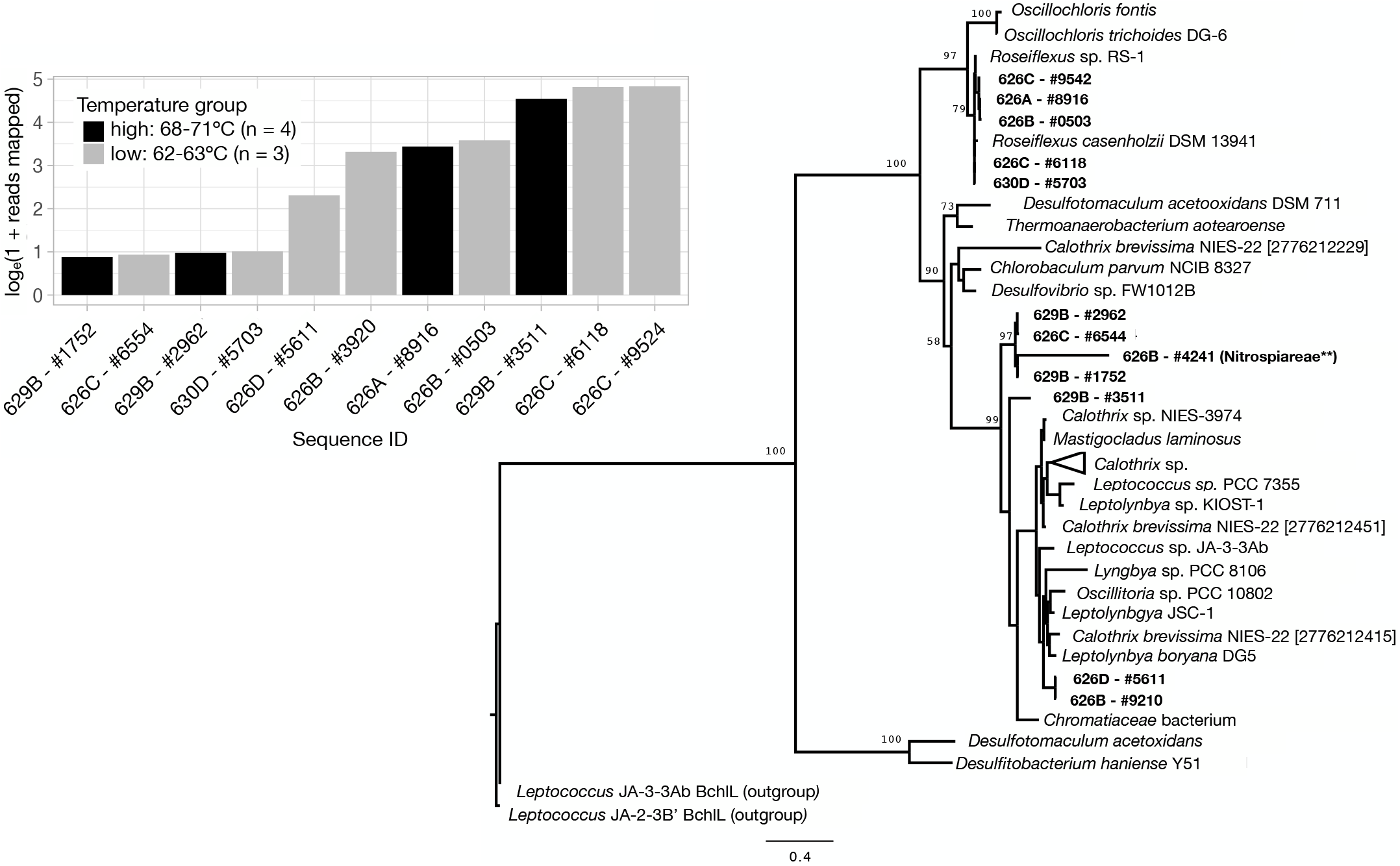
Maximum likelihood-inferred NifH tree. Maximum likelihood-inferred phylogenetic reconstruction of NifH sequences (12) recovered from YNP sites (bold) aligned to 22 references sequences. The tree is rooted with Leptococcus BchlL. The scale bar indicates 4 substitutions for every ten positions. Bootstrap values (n=1000) are displayed at branches. Bootstrap values less than 50 are not shown. Overlain the tree is bar plot displaying the natural log(1 + reads mapped) of each NifH in the tree, shaded by high (68-71°C) and low (62-63°C) temperature group. ** Indicates the result of BLASTP search.

Finally, the third most abundant *nifH* in our dataset (629B #3511) formed a separate clade near, but not within, the cyanobacteria clade (Figure 4). Protein BLAST analysis (27) revealed that these NifH sequences are Aquificae-like NifH, a deep-branching chemolithoautotrophic group with diazotrophic representatives found in high temperature (>70°C) hot springs (17). Previous analysis of *nifH* genes across all domains of life suggested Aquificae are the oldest extant diazotrophic bacteria (19). Thus, our data contain several *nifH-*containing lineages that are of great importance for solving the evolutionary history of nitrogen fixation.

### Metagenome assembled genomes reveal distinct differences between Chloroflexus and Roseiflexus species

Phototrophic Chloroflexi species are widely distributed and abundant in alkaline hot springs > 60°C, but their contribution to carbon and nitrogen cycling in hot springs is not well understood. Through metagenome assembly data, we have begun to piece together the metabolic potential of phototrophic Chloroflexi *in situ*. We have shown that genes for phototrophic machinery in cyanobacteria and Chloroflexi are abundant throughout the eight hot spring samples surveyed here. Key genes in photoautotrophic Chloroflexi (3HPB pathway) genomes are variable with temperature (*mcr* is less abundant in low temperature sites, Figure 3D), but generally abundant across sites. Additionally, a large proportion of the NifH sequences we recovered were most closely related to *Roseiflexus* species. To examine the distribution of specific Chloroflexi taxa, we binned our metagenome assemblies and quantified the abundance of bins classified as Chloroflexi taxa in each site (Figure 5A, S5, Table S2). Furthermore, we conducted a BLASTN search (27) of the Chloroflexi metagenome bins for genes involved in carbon fixation, Type-II reaction center photosynthesis and NifH/BchL protein families (Figure 5B).

**Figure 5.**
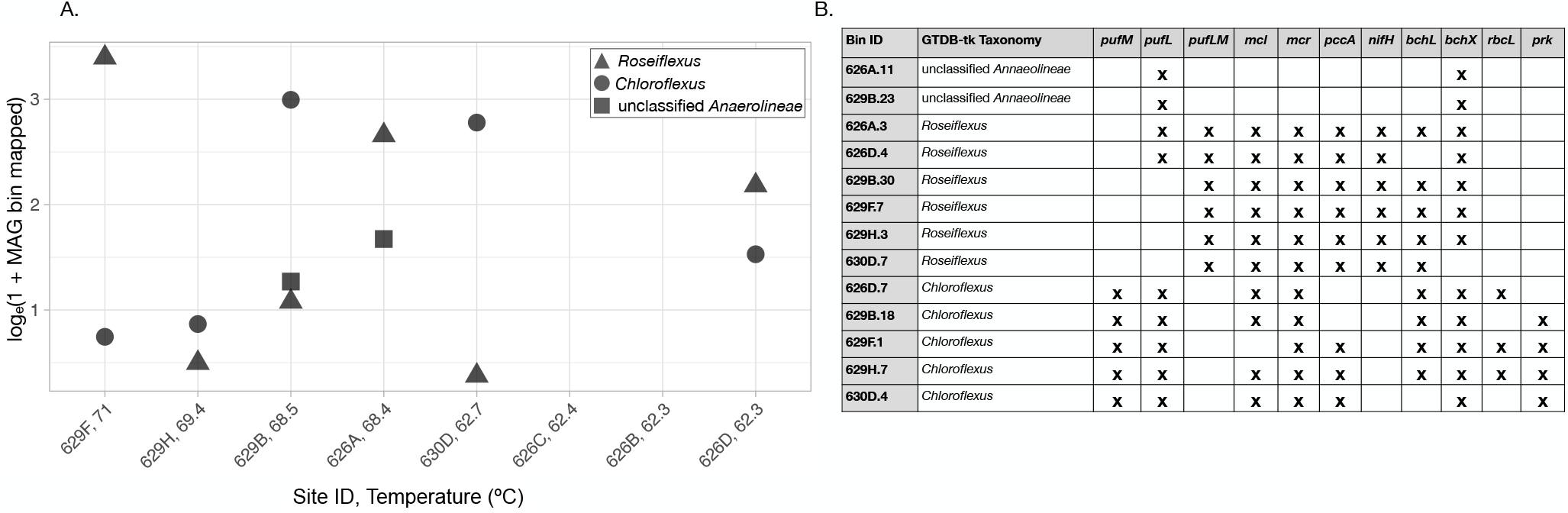
Phototrophic Chloroflexi bins have distinct distributions with temperature. Metagenome bins were mapped to assemblies to determine the proportion of the metagenome accounted for by that bin and natural log (1 + reads mapped) transformed. Each point represents an individual MAG (see Table S2). MAGs are plotted for each site and sites are ordered by decreasing temperature. GTDB-tk assigned taxonomy is differentiated by shape. (B) BLASTN hits for select genes in the metagenome bin. Note: *rbcS* was not recovered in any of the bins and is, thus, excluded from the table.

We recovered two putative, phototrophic Chloroflexi genera from our binning analysis: *Roseiflexus* and *Chloroflexus.* These two genera were represented by several distinct bins across multiple sites (Figure 5A, Table S2). *Roseiflexus* and *Chloroflexus* bins were more abundant than non-phototrophic Chloroflexi, consistent with our gene analysis above and previous work in alkaline hot springs (8,9,12). *Roseiflexus* bins comprise a large proportion of the metagenome in several sites (Figure 4A), while *Chloroflexus* bins were generally more abundant in sites > 68°C. Furthermore, *Roseiflexus* bins appear to be functionally different than *Chloroflexus* bins in our metagenomes (Figure 5B). Each *Roseiflexus* bin contained fused *pufLM* sequences, all three 3HPB genes from our assembly analysis (*mcl, mcr* and *pccA)* and *nifH* sequences. Not surprisingly, none of our *Roseiflexus* bins contained genes involved in the CBB cycle, which has not been found in any of the sequenced *Roseiflexus* genomes (50,52). Since *Rosieflexus* are dispersed throughout these sites (12), abundant in our metagenomes (Figure 5A) and encode the 3HPB pathway, it is likely that they are living autotrophically or mixotrophically *in situ*.

Alternatively, several of our *Chloroflexus* bins harbored a mosaic of genes involved in photoautotrophy. Each *Chloroflexus* bin contained *pufL* and *pufM* genes, which would support the characteristic phototrophic lifestyle of this group (54). However, three of the five *Chloroflexus* bins were missing genes that are critical to the 3HPB pathway, but the bins contained *prk* or *rbcL* genes (Figure 4B). This suggests that a handful of our *Chloroflexus* may have lost a functional 3HPB pathway, but are instead operating a modified CBB cycle to fix carbon, which has been suggested in other systems (55). Previous studies have implied that photoautotrophy is a characteristic of *Chloroflexus*, while *Roseiflexus* are photoheterotrophs (6,14,50,56), but our metagenome bin analysis coupled with ^13^C fractionation suggest *Roseiflexus* could be contributing to carbon fixation *in situ* and that *Chloroflexus* strains could be using the CBB cycle to fix carbon. Lastly, nitrogen fixation and bacteriochlorophyll synthesis genes share an evolutionary history (57); therefore, we would expect that closely related species like *Roseiflexus* and *Chloroflexus* would both contain these genes. Yet, our data suggest that only *Roseiflexus* encode both *nifH* and *bchL,* while *Chloroflexus* contain only *bchL* genes, which is in accordance with previous work (50,52,53).

While phototrophy in phylum Chloroflexi is thought to be limited to the class *Chloroflexales*, a draft genome classified within the candidate genus *Roseilinea,* was recovered from a sulfidic spring and appears to live a photoheterotrophic lifestyle (13). *Roseilinea* is within the non-phototrophic class Anaerolineae and is only distantly related to *Chloroflexus* and *Roseiflexus* species. Two of our metagenome bins were classified as ‘Anaerolineae unclassified’ (626A.11 and 626B.23) and both contain *pufL* and *bchX* sequences, suggesting they are likely within the novel *Roseilinea* genus. Both of these alkaline sites were low in sulfide, which differs from the acidic and sulfidic Japanese hot springs where the *Ca.* Roseilinea mizusawaensis AA3_104 genome was recovered (13). Additionally, our metagenomes contained several unclassified or novel Chloroflexi bins (Figure 4, S6, Table S2), suggesting high temperature alkaline, hot springs are a suitable environment to tease apart the physiology of novel, phototrophic Chloroflexi *in situ.*

In summary, our metagenome data indicate a decrease in the diversity but not necessarily in the abundance of photoautotrophs with increasing temperature: genes associated with cyanobacteria (*psb* and CBB pathway genes) were more diverse at lower temperatures, but abundant across sites while key genes in the 3HPB pathway are present between 60-71°C Our data are consistent with high temperature cyanobacteria are highly adapted to specific temperatures in alkaline hot springs and suggest temperature selects for specific Chloroflexi taxa as well: *Chloroflexus* bins were most abundant in sites below 63°C while *Roseiflexus* bins that contain genes necessary for the 3HPB pathway were abundant in sites with varying temperatures. Furthermore, *nifH* genes were abundant across sites, regardless of site temperature and *Roseiflexus*-like NifH were among the most abundant in our data. While *Roseiflexus* are likely not fixing nitrogen *in situ*, *nifH*-containing *Roseiflexus* could be critical to solving the evolutionary puzzle of nitrogen fixation in bacteria. While previous work has suggested that *Roseiflexus* and *Chloroflexus* are heterotrophs and autotrophs, respectively, our work suggests that both taxa could live a mixotrophic lifestyle *in situ*.

## Materials and Methods

### Data collection, geochemistry and sample processing

Biomass from eight sites in YNP (Table 1) were collected and processed as previously described (10). Briefly, samples were collected in 2017 using sterilized forceps or pliers and stored on dry ice in transit. DNA (~250mg) was extracted using the QIAgen powersoil kit following the manufacturer’s protocol. Sulfide, Fe^2+^, and dissolved silica were measured onsite using a DR1900 portable spectrophotometer (Hach Company, Loveland, CO). Water samples were filtered through 0.2-μm polyethersulfone syringe filters (VWR International, Radnor, PA, USA) and analyzed for dissolved inorganic carbon (DIC) concentration, δ^13^C and δ^15^N as described previously (20). Field blanks comprised of filtered 18.2 MΩ/cm deionized water, transported to the field in 1-liter Nalgene bottles (acid washed as described above), were collected onsite using the equipment and techniques described above.

### Metagenome sequencing, assembly and binning

Total DNA for eight samples was submitted to the University of Minnesota Genomics Center (St. Paul, Minnesota, UMGC) for metagenomic sequencing with an Illumina HiSeq 2500. The UMGC prepared dual indexed Nextera XT DNA libraries following the manufacturer’s instructions for each sample. The samples were sequenced on two lanes, generating >220M 1×125 bp reads. The mean quality scores were >Q30 for all libraries. Reads were trimmed using Sickle (v. 1.33) with a PHRED SCORE > 20 and a minimum length threshold of 50 (58) and assembled using SPades (v. 3.11.0) (59) using the default parameters. Initial binning of metagenome assemblies was conducted using Metabat2 (v.2.12.1)(60) and assessed for completeness and contamination via CheckM (v. 1.1.3) (61). Bins were considered clean if they were <10% contaminated and > 70% complete – bins that did not meet this threshold were further refined using DASTool (v. 1.1.2)(62). DASTool takes bins from multiple binning algorithms and creates a set of ‘best’ bins. To run DASTool, we re-binned assemblies containing ‘unclean’ bins with MaxBin (v. 1.2.9) (63) and CONCOCT (v 1.0.0)(64), in addition to MetaBat, and input the bins from three binning algorithms into DASTool following the default settings. DASTool bins were assess for quality via CheckM. Quality-filtered sequence reads mapped to each bin in a sample by the program BBMap (65) were used to calculate percent mapped reads as a proxy for bin relative abundance. All bins were assigned taxonomy using GTDB-tk (v 1.3.0) (66). Final bin classification and statistics are displayed in Table S2. A BLASTN search using the default settings (27) for photosynthesis and *nifH* gene families was conducted genes from isolate genomes (see Supplemental material).

### Annotation and comparison of functional genes

Metagenome assemblies for eight sites (Table 1, Table S1) were submitted to the Joint Genome Institute for structural and functional annotation via the DOE-JGI Microbial Genome Annotation Pipeline. Briefly, open reading frames (ORFs) were predicted using Prodigal (67) and the resulting amino acid sequences were assigned functional annotations. Select genes (see supplemental files for complete list and KEGG IDs) involved in three carbon fixation pathways (the Calvin Cycle, 3-Hydroxypriopionate Bicycle, and the reverse Tricarboxylic Acid cycle), nitrogen fixation and photosynthesis were queried in the annotated assemblies. Genes of interest were retrieved using search tools on the Joint Genome Institute web interface and known KEGG Orthologies. Metagenome reads were mapped to each JGI ORF using Bowtie2 (68). All reads were mapped at > 90% alignment rate.

To determine abundance of select genes involved in photosynthesis, carbon fixation, and nitrogen fixation, number of reads mapped to genes of interest were calculated using the pileup.sh script in BBTools (65). In order to directly compare genes of interest, genes were normalized by gene length and metagenome size using the following equation:

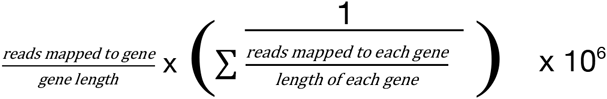

If multiple ORFs were assigned to a functional annotation, the normalized read abundance for that functional annotation was averaged. All analysis of functional genes, plotting and statistical analysis was conducted in R v3.6.1(69) using the following packages: *tidyverse* (70), *ggplot2* (71) and *vegan* (72). All metagenome assemblies are available on JGI (see Table 1, Table S1).

### Phylogenomic tree construction

Phylogenetic trees for both NifH and PsbA were built by compiling the proteins sequences of interest from the functionally annotated metagenome assemblies with reference protein sequences. Reference sequences were obtained on UniPort (73). Briefly, sequences < 200 amino acids were removed, sequences were aligned with MUSCLE v3.8.31(default parameters) (74) and alignments were trimmed using Gblocks (default parameters with the exception of *-b5-h*) (75). Phylogenetic analysis with bootstrap support (n=1000) of trimmed, aligned protein sequences was conducted using RAxML (v. 8.2.11) (76) using the PROTGAMMAJTTF substitution model. The subsequent newick file was edited using FigTree (v. 1.4.4) (77) to generate trees.

## Acknowledgements

T.L.H. conducts research in Yellowstone National Park under research permit YELL-2018-SCI-7020 issued by the Yellowstone Research Permit Office and reviewed annually. This work was supported by the University of Minnesota. The authors acknowledge the Minnesota Supercomputing Institute (MSI) at the University of Minnesota for providing resources that contributed to the research results reported within this paper. We are grateful to the entire staff of the Yellowstone Research Permit Office for facilitating the permitting process to perform research in YNP. Special thanks to Annie Carlson and Erik Oberg in the Yellowstone Research Permit Office. We would like to thank J. Havig, L. Brengman, C. Grettenberger, L. Seyler, and J. Kuether for technical assistance in the field, A. Borowski and K. Quinn for assistance processing samples in the lab and H. Sauer for troubleshooting statistical analyses.

## AUTHOR CONTRIBUTIONS

A.C.B and T.L.H. designed the study and completed the laboratory analyses. A.C.B. and T.L.H. collected samples and performed the field work. A.C.B, S.K.M, E.K. and T.L.H. analyzed the data. A.C.B interpreted the data and wrote the manuscript with contributions from S.K.M, E.K. and T.L.H.

## MATERIALS & CORRESPONDENCE

Correspondence and requests for materials should be addressed to T.L.H.: Trinity L. Hamilton. Department of Plant and Microbial Biology, University of Minnesota, St. Paul, USA, 55108. Phone: +16126256372

## SUPPLEMENTAL FIGURE LEGENDS

Table S1. Metagenome assembly statistics.

Figure S1. Type I reaction center genes.

The overall abundance (natural log of 1 + reads mapped) of type I anoxygenic photosynthesis reaction centers are shown as box plots for each site. Triangles represent the mean abundance for the gene set and dots represent individual gene abundances, shaded by reaction center gene. Box plot outliers were removed. Sites are ordered by decreasing temperature.

Figure S2. Ratio of CBB genes with temperature.

The ratio of *rbcS* genes (x axis) to both *rbcL* and *prk* genes (y axis) is shown. Points are shaded by site temperature.

Figure S3. rTCA cycle gene distribution with temperature.

The mean abundance (natural log of 1 + reads mapped) for the rTCA cycle plotted as boxplots for each site. Triangles represent the mean abundance for the gene set and dots represent individual gene abundances, shaded by gene. Box plot outliers were removed. Sites are ordered by decreasing temperature. Kruskal-Wallis H test significance values (p-value) for each gene are displayed to the right of the boxplot.

Figure S4. Distribution of *nifH* and alpha diversity of genes with temperature.

(A) Natural log (1 + reads mapped) for *nifH* genes recovered from each site, ordered by decreasing temperature displayed as boxplots. Triangles represent the mean. (B) Distinct *nifH* genes recovered from each site, ordered by decreasing temperature. Dashed vertical line separates sites > 68°C and < 63°C. (C) Spearman-rank correlations for *nifH* natural log (1 + reads mapped) for temperature, molybdenum, sulfide and total iron in each site. Shaded area represents the standard error, R^2^ and p-values are displayed in the upper right corner of each plot.

Figure S5. Abundance and alpha diversity of all Chloroflexi bins.

Metagenome assembled genomes (MAGs) were mapped to assemblies to determine the proportion of the metagenome accounted for by that bin and natural log (1 + reads mapped) transformed for all Chloroflexi bins. Phototrophic Chloroflexi includes class *Chloroflexales* and Non-Phototrophic Chloroflexi includes the remaining classes. Each point represents an individual MAG (see Table S2). MAGs are plotted for each site and sites are ordered by decreasing temperature. GTDB-tk assigned taxonomy (“Genus”) is differentiated by shaded points. (B) Number of distinct Chloroflexi bins in each site is plotted, ordered by temperature.

Table S2. Chloroflexi bins statistics determined by CheckM and GTDB-tk.

